# Sex Differences in Head-fixed Running Behavior

**DOI:** 10.1101/585000

**Authors:** Emily J. Warner, Krishnan Padmanabhan

**Affiliations:** Department of Neuroscience, University of Rochester School of Medicine and Dentistry, Rochester NY 14642; Neuroscience Graduate Program (NGP), University of Rochester School of Medicine and Dentistry, Rochester NY 14642; The Ernest J. Del Monte Institute for Neuroscience, University of Rochester School of Medicine and Dentistry, Rochester NY 14642; Center for Visual Science, University of Rochester School of Medicine and Dentistry, Rochester NY 14642

## Abstract

Sex differences in running behaviors between male and female mice occur naturally in the wild. Recent experiments using head restrained mice on a running wheel have exploited locomotion to provide insight in the neural underpinnings of a number of behaviors ranging from spatial navigation to decision making. However, it is largely unknown how males and females behave differently in this experimental paradigm. We found that in head-fixed mice that were initially exposed to a running wheel, all female mice ran forward naturally within the first two days, while almost all male mice scurried backward for up to 4 days. With daily exposure, male mice progressively learned to naturally run forward, with this transition occurring over the course of a 7-day period. Taken together, we have identified a sexually divergent behavior in head-fixed running that should be considered in experiments that use this experimental design. Furthermore, this sex-specific difference could serve as a new way to interrogate the neural underpinnings of a number of behaviors such as anxiety or fear.

## Introduction

Across multiple species, variability in behavior due to sex differences can be traced to differences in neural circuits (Mowrey & Portman, 2012; Yang & Shah, 2016). In mice, for instance, behaviors as diverse as fear conditioning and navigation on the Morris water maze vary based on the sex of the animal (Roof & Stein, 1999; Keeley *et al.*, 2013; Yang *et al.*, 2013; Gruene, Flick, *et al.*, 2015). Examples of sex-specific differences in behavior can be found outside of the domain of fear and learning. For instance, more prosaic behaviors, such as the distance or the duration that an animal runs vary between males and females in the wild (Lightfoot *et al.*, 2004; Goh & Ladiges, 2015). Thus, although a number of behaviors studied in laboratory settings may be sexually divergent, much of what is known about behavior either uses exclusively male mice (Tronson, 2018) or may not treat sex as a dependent variable when using male and female animals (Shansky & Woolley, 2016). For example, a common experiment involves head-fixing an awake behaving rodent and placing it onto a running wheel to study the circuits involved in sensory processing (Niell *et al.*, 2010; Smear *et al.*, 2011) spatial navigation (Harvey *et al.*, 2009; Dombeck *et al.*, 2010), and decision making (Abraham *et al.*, 2010; Smear *et al.*, 2013; Juavinett *et al.*, 2018). Despite the ubiquity of this paradigm in systems neuroscience, and the importance of measuring running either as a feature, or a confound of experiments, it remains unclear if there are differences between males and females in head-restrained running.

To explore this, we analyzed the running behavior of male and female adult mice over a seven-day period while head-fixed on a running wheel. First, in both males and females, we found an increase in the probability of running and the velocity with which animals ran over the 7-day period. Most interestingly, however, we saw significant sex differences in the direction that mice ran during the early days of exposure to the run-wheel. Within 2 days on the running wheel, all female mice ran forward, while male mice scurried backward. Over multiple days. It was not until 4-5 days of exposure in a 7-day training period that male mice began to run naturally. Thus, by studying not only the speed but the velocity of running, we unmasked a sex-specific divergent behavior in mice acclimating to a common laboratory experimental paradigm.

## Materials and Methods

### Animals

18 mice, 9 male and 9 female C57BL/6 mice, 3-4 months old were utilized for this experiment. All experiments were performed in accordance with the guidelines for animal use and care as outlined by the University Committee on Animal Resources (UCAR) at the University of Rochester Medical Center.

### Head-Fix Procedure

Prior to procedure, animals were dosed with 0.5-1.0mg/kg slow-release buprenorphine via subcutaneous injection. Animals were anesthetized using inhalation of between 1-2% vaporized isoflurane and then placed in a stereotaxic for surgery. Following a midline incision on the skull connective tissue was resected and excess skin removed and vetbond was placed to attach the perimeter skin to the skull. A 3D printed head frame was then put into place and dental cemented to the base of the skull taking care to provide enough clearance for the ears. The area was then allowed to dry completely prior to placing the animal into the home cage for recovery. Animals were recovered for 24 hours prior to behavioral habituation.

### Running Wheel Habituation

Animals were trained on the running wheel beginning 24 hours after head-fix procedure for seven consecutive days. Mice were weighed daily prior to habituation to ensure that animals were not losing significant body weight and to confirm that the running behavior was not affected by the head-fixing procedure itself. Animals were habituated for one hour per day on a circular running wheel that allowed for both forward and reverse running, with the last fifteen minutes of run behavior recorded. While animals were monitored remotely with a camera, all habituation took place in darkness, and during habituation recording, there was no intervention or light input. Habituation was consistently completed during the animals’ light cycle within the vivarium.

## Results

To explore the question of sex differences in head-fixed animals, we first implanted a 3D-printed interface to the animals’ skull to allow the animal to be stabilized on a run-wheel. Following this, animals were allowed 24 hours to recover from surgery prior to 7 consecutive days on a run-wheel. Beginning on day 1, mice were placed on a one-dimensional running wheel with their head fixed for 1 hour per day (**Fig. 1A**), allowing them to rotate the wheel forward, backward, or remain stationary. Importantly, in this paradigm, we could evaluate differences between forward running, which was most analogous to natural running in mice, and reverse running, which mimics a scurrying or escape behavior, most analogous to anxiety in mice (Gogas *et al.*, 2007). A 15 trace from the 1 hour period (**Fig. 1B**) for a single animal on day 1 showed epochs of running in the forward direction (positive velocity in the gray region), and in the reverse direction (negative velocity in the white region, **Fig 1B**). Interspersed between the bouts of running were periods where the animal remained stationary. To quantify these features of running, we calculated the percentage of time the animal was running, as opposed to remaining stationary. Across all animals, there was a significant increase in the percent that an animal was running between day 1 and day 7 (n=18; day 1, 0.2147; day 7, 0.4224; p = 0.0016; Wilcoxon rank-sum test; **Fig 1C**). Furthermore, we found a significant increase in the overall velocity of running from day 1 to day 7 (n=18; day 1, 0.8008 cm/sec; day 7, 3.9380 cm/sec; p = 0.00007; Wilcoxon rank-sum test, **Fig. 1D**). Taken together, over the 7-day period as the animals grew accustomed to the run-wheel, they ran faster and more frequently. To determine if these differences persisted after 7 days, we continued to measure the running behavior in animals from greater than or equal to 20 days (N=2) and found no significant increase in run velocity or run duration (data not shown) over this prolonger period. Thus behavior settled after 7 days on the run wheel.

**Figure 1.**
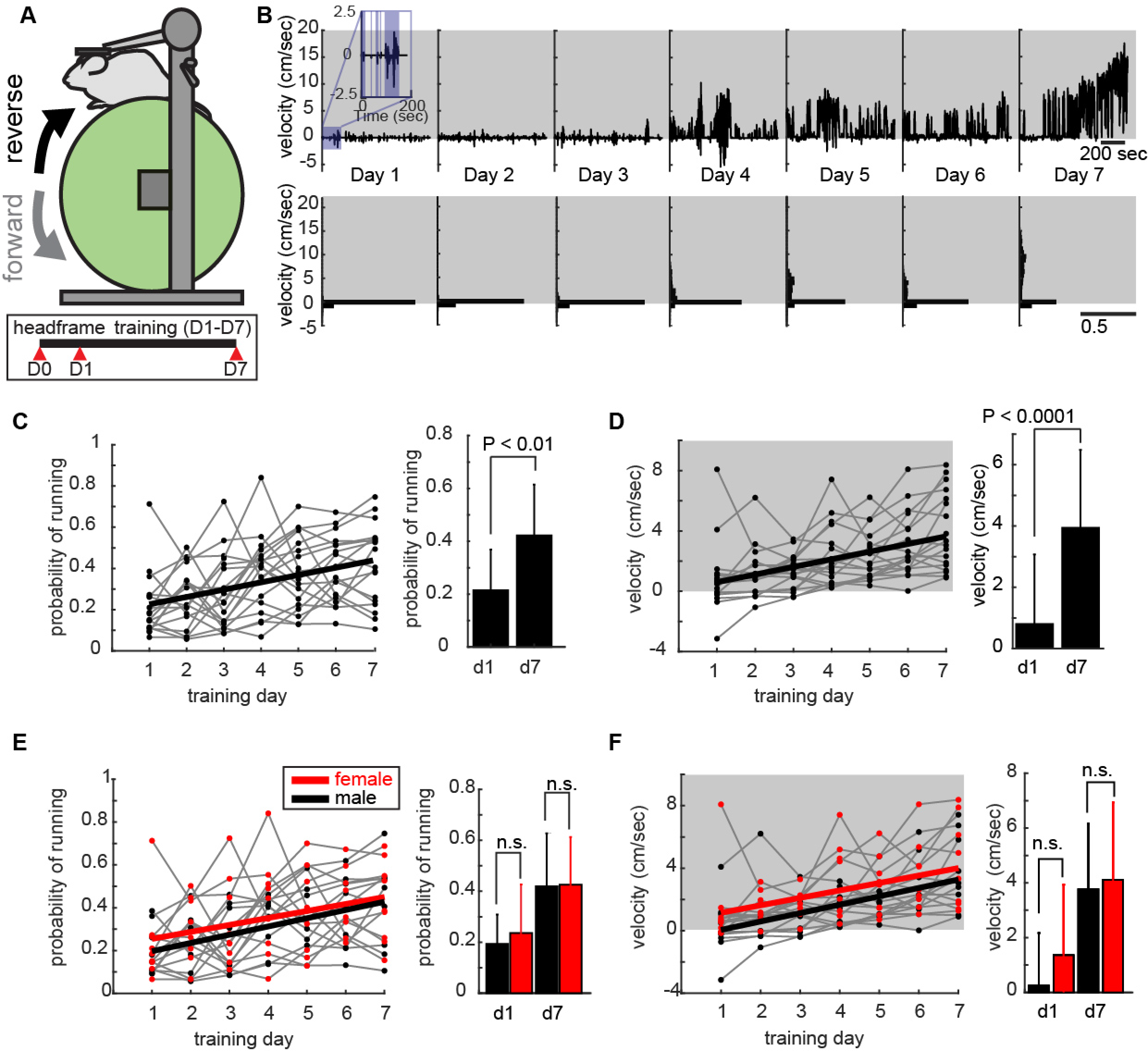
The velocity and distance covered increases over a 7-day period. (A) Voluntary running wheel schematic where head-fixed mouse can freely run on a 1-dimensional run-wheel. (B) Example running behavior trace where shaded gray area represents running in the forward direction and non-shaded area is running in the reverse direction. Inset: blue shaded areas show epochs of running and non-shaded shows stationary epochs where the animal is not running. C) Mean probability of running for all animals (n=18). Left: Gray lines are for each individual animal; solid black line shows mean probability of running. Right: quantification of probability of running for all animals from day 1 and day 7 of habituation. Error bars are standard deviation, Wilcoxon rank sum test. (D) Velocity analysis for all animals (n=18). Left: Shaded area represents running in the forward direction. Gray traces are for individual animals, solid black line for mean velocity over the 7 days of training for all animals. Right: quantification of velocity for all animals between day 1 and day 7. Error bars are standard deviation, statistical test used: Wilcoxon rank sum test. (E) Probability of running for males and females. Left: data from C separated into males (black) and females (red). Right: quantification of probability of running between males and females on day 1 and day 7 of habituation. Error bars are standard deviation, statistical test used: Wilcoxon rank sum test. (F) Velocity for males and females. Left: The data is from D separated into males (black) and females (red). Right: quantification of velocity between males and females on day 1 and day 7 of habituation. Error bars are standard deviation, statistical test used: Wilcoxon rank sum test.

Although there were general trends in the running over the first 7 days of habituation, we noted that there was a large spread in the behavior, particularly over the first three days. For example, the standard deviation of the speed when then animals ran (D1 = 2.2762 cm/sec) was much greater during the first day as compared to the third day of habituation (D3 = 1.1510 cm/sec, F-statistic, p=0.0075) suggesting that there could be important inter-individual differences when the animals were first acclimating to the run-wheel.

To determine if sex differences could account for some of this variability, both the probability of running and speed of running were analyzed for males versus females separately (**Fig. 1E, F**). Although there were no significant differences between males and females on day 1 and day 7 for both probability of running and speed, we did see a trend where females ran more often and faster than males. To further explore these differences, we dissected the specific aspects of running behavior across male and female animals.

As animals could run either forward or reverse on the wheel (**Fig. 1B**), we wished to determine if the trends in running between males and females could be accounted for not only by considering the magnitude (speed), but also the direction (velocity) of running (**Fig. 2A**). To do this, we plotted the cumulative sum of the distance the animal ran over the training period and dividing it by the final position of the animal to calculate a normalized position of running. If the animal ran mostly forward, then the net normalized position at the end of habituation would be +1. Conversely, if the animal ran largely in the reverse direction, its net normalized position would be −1. Across all animals, on the first day, we saw a wide distribution of running in the forward and reverse direction with both males (**Fig 2A, black**) and females (**Fig. 2A, red**) as evidenced by the range in the net normalized position. By day 2 however, we began to see distinct differences in running emerge between males and females. On this day, almost all the females ran in the forward direction (N=8/9) while the majority of males continued to scurry in the reverse direction (N=5/9). This difference persisted into day 3, and it was only by day 4 that all the males began running forward.

**Figure 2.**
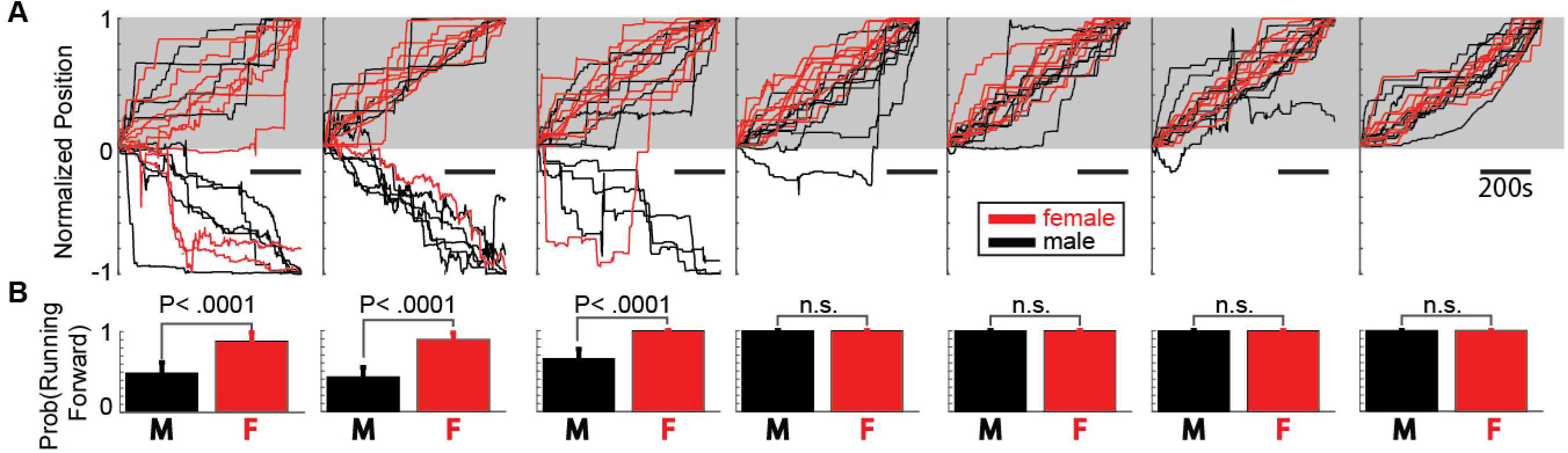
Female mice run forward earlier in 7-day period as compared to male mice. (A) Normalized position of running behavior. Net forward running (in gray shaded region) results in an end position at +1. Net reverse running (non-shaded region) results in end position at −1. (Scale bar = 200 s) (B) Quantification of probability of running forward versus reverse for males and females across days of habituation (n=18). Error bars are standard deviation, statistical test used: Wilcoxon rank sum test and bootstrap with replacement.

To further quantify differences in the direction of running, we measured the probability that animals within a group run forward on any given day (**Fig. 2B**). During the first day on the run-wheel, the probability of running forward was significantly higher for females than it was for males (n=9 males, n=9 females; day 1 male, 0.4333 ± 0.1185; day 1 female, 0.7733 ± 0.0945; p=<0.0001; Confidence intervals estimated by Bootstrap with replacement, Wilcoxon rank-sum test). This difference remained significant for the second (n=9 males, n=9 females; day 2 males, 0.4333 ± 0.1185; day 2 females, 0.8889 ± 0.0735; p=<0.0001, Wilcoxon rank-sum test) and third day on the run wheel (n=9 males, n=9 females; day 3 males=0.6422 ± 0.1126; day 3 females=1; p=<0.0001; Wilcoxon rank-sum test). By day 4, and for the duration of days the mice were exposed to the run wheel, all male mice ran forward; any sex-specific differences in head-fixed running behaviors were consequently abolished.

In addition to the differences we observed between males and females, we wished to identify in any of the specific features of the behavior could provide insight into the transition from scurrying to running, and capture the diversity of strategies employed across days by each animal. A plot of normalized position for two males (blue = day 1, green = day 7) and two females (red = day 1, yellow = day 7) over the seven days revealed three hallmarks in the behavior that illustrated key features of running (**Fig. 3A**). First, on the days when the animals switched from scurrying to running forward, the moment of this transition was abrupt, and almost always included a prolonged bout of rapid running in the forward direction. We observed this transition in 7/9 males and 3/9 females, although the day on which this transition occurred varied across individuals and across the two sexes. Second, we observed that once animals ran in a forward direction, they all exhibited a stereotypic pattern of locomotion, running forward for brief epochs followed by periods where they remained stationary as illustrated by plots of the acceleration for the 4 example animals across days 5-7. In each case, periods of rapid forward movement (**Fig. 3B, black arrows**) were interleaved with periods where the animal was stationary (**Fig. 3B, gray arrows**). These data show that although each individual covered a different distance over the 1-hour period, the way in which they cover that distance (periods of running and stationary) was common across all individuals. Finally, we found that once an animal transitioned from scurrying backward on the wheel to running forward, they did not revert their behavior (**Fig. 3C**). We represented this as a change matrix for all the males and females across the 7 days, where a white box was a change from running in the reverse direction on the previous day to running in the forward direction, a black box indicated no change in direction of running and a gray box corresponded to a change running from the forward direction to the reverse direction. Importantly, we saw no gray boxes, showing that once a transition occurred from backward scurrying to forward running, this transition was permanent.

**Figure 3.**
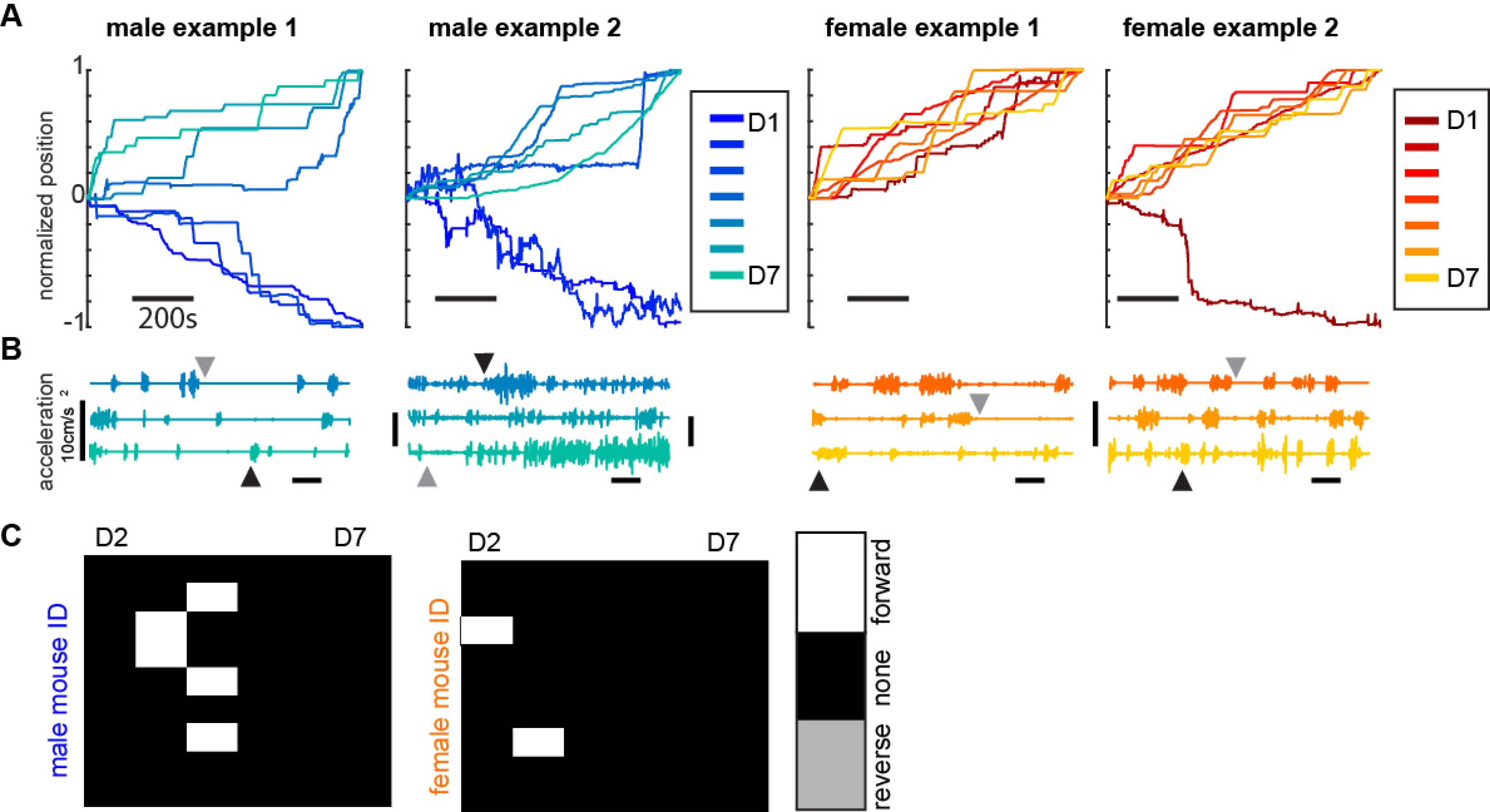
Inter-individual differences in head-fixed running behavior. (A) Normalized position for male (n=2) (blue) and female (n=2) (red) examples. Running in the forward direction results in an end position at +1 while net reverse running results in an end position at −1. (B) acceleration plots for males and females on the last three days of habituation (day 5-7). Black arrows indicate active running while gray arrows indicate periods where the mouse is stationary. (D) Differential plots. Columns represent the difference between the day listed and the day before. Rows are individual animals. White box shows change in running direction from reverse to forward running. Black box is no change in direction. Gray is a reversion from running forward to running in reverse.

## Discussion

In this work, we identified sex differences in running behavior of head-fixed mice over a 7 day exposure period with the females learning to run forward before the males. Although this difference is apparent initially, by day 4 of habituation all the animals, both male and female, run in the forward direction. Additionally, while we found that on average female mice ran farther and more frequently than male mice, this difference was not statistically significant. On the individual level, there are several hallmarks of running behavior that are seen in every animal, regardless of sex. These include a sudden jump in running activity when the animal begins to run forward, dynamics the include epochs of running and periods of remaining stationary, and never reversing direction after learning to run forward. Our findings suggest that sex differences in natural run behavior (Lightfoot *et al.*, 2004; Goh & Ladiges, 2015) are recapitulated in head-fixed laboratory experiments. Importantly, these sex differences emerge only when the behavior of animals is dissected in detail, for instance by studying not just the amount that an animal runs (speed), but also identifying if that running happens in the forward direction or the reverse scurrying direction (velocity).

A number of recent studies highlight the importance of considering sex as a dependent variable when analyzing behavior in the laboratory (Lin *et al.*, 2011; Yang *et al.*, 2013; Gruene, Flick, *et al.*, 2015; Gruene, Roberts, *et al.*, 2015; Kim *et al.*, 2015). For instance, sexually divergent responses emerge when Pavlovian fear responses in classical conditioning experiments are analyzed not only in terms of whether an animal freezes or not, but the *kind* of motion that happens if an animal does move (Gruene, Flick, *et al.*, 2015). Not unlike these experiments, the initial introduction to the head-fixed run wheel could represent a novel aversive environment that the mouse is accustomed to with increasing days of experience. It is therefore not surprising that sexually divergent strategies for running would emerge over the initial habituation period. Consequently, in experiments ranging from those involving spatial navigation (Dombeck *et al.*, 2010; Meshulam *et al.*, 2017), to those that study running modulation of sensory coding (Niell *et al.*, 2010), the consideration of sex could be paramount in interpreting results (Shansky & Woolley, 2016). Additionally, it is prudent in studies that use head-fixed behaviors to monitor the details of the behavior during the early phases of habituation as these differences may impact the interpretation of results later in the experiment. Furthermore, our data suggest that the early differences in head-fixed running could be a novel experimental framework to investigate the natural diversity of neural circuits involved in fear (Pibiri *et al.*, 2008; Hauner *et al.*, 2013; Yang & Shah, 2016), spatial reasoning (Harvey *et al.*, 2009), and anxiety (Zeng *et al.*, 2011; Ciocchi *et al.*, 2015). Finally, beyond their relevance to understanding the natural diversity of behaviors (Shansky & Woolley, 2016) the inclusion of female animals in experiments can provide insight into divergent circuits that shape these behaviors (Yang & Shah, 2016; Tronson, 2018) and the extent to which such differences translate to different vulnerabilities to neurological and psychiatric disorders based on sex (Earls, 1987; The Lancet Neurology, 2019).

## Acknowledgments

KP supervised this project. EJW performed the experiments. EJW and KP performed the analysis and wrote the manuscript. KP is supported by NIMH R00 MH101634, NIMH R01 MH113924, NSF CAREER Award, the Cystinosis Research Foundation, and the Schmitt Foundation.

